# Deep Learning-Based Genetic Perturbation Models *Do* Outperform Uninformative Baselines on Well-Calibrated Metrics

**DOI:** 10.1101/2025.10.20.683304

**Authors:** Henry E. Miller, Gabriel M. Mejia, Francis J. A. Leblanc, Bo Wang, Brendan Swain, Lucas Paulo de Lima Camillo

## Abstract

Single cell genetic perturbation modeling involves predicting the effects of unobserved genetic manipulations, enabling scalable *in silico* screens for target discovery. Recent reports have claimed that deep learning-based perturbation models fail to outperform uninformative baselines, raising doubts about their utility. Here, we show that these conclusions largely stem from limitations of benchmarking *metrics*, not from the models themselves. We introduce a framework for evaluating benchmark metric calibration using positive and negative controls, including a new positive control baseline (the *interpolated duplicate*) and a quantitative calibration measure (the *dynamic range fraction*). Across 14 perturbation datasets and 13 evaluation metrics, we find that conventional metrics such as mean squared error (MSE) and control-referenced delta correlation (Pearson(Δ_ctrl_)) are often poorly calibrated, whereas weighted and rank-based alternatives exhibit consistent calibration. Under well-calibrated metrics, deep learning models outperform mean, control, and linear baselines, and in some cases even surpass the additive baseline in combination-prediction tasks. Calibrated evaluation thus explains prior reports of model underperformance, revealing that deep learning models *do* outperform uninformative baselines.

## Introduction

Single-cell genetic perturbation modeling aims to predict transcriptomic responses to gene perturbations (typically inhibition or activation) in one or more biological contexts (e.g., cell types). If successful, perturbation response models (sometimes termed “virtual cells”^1^) could enable scalable *in silico* screens to uncover genes which reverse disease or restore youthful cellular states. Recent studies, however, have questioned whether transcriptome-level perturbation modeling is viable. In their recent work, Ahlmann-Eltze et al. (2) reported that deep learning models do not outperform uninformative baselines across multiple datasets. Likewise, several independent benchmarks have found that a simple “mean baseline,” the average perturbation response in the training set, often matches or surpasses state-of-the-art models on metrics such as mean absolute error (MAE), mean squared error (MSE), or the correlation of control-referenced deltas (Pearson(Δ_ctrl_)) (3–8). Together, these findings invite skepticism about the utility of deep learning-based perturbation response models.

In earlier work, we identified two artifacts that explain why the mean baseline can appear competitive despite lacking perturbation-specific signal (9). The first, *control bias*, arises when systematic differences between control and perturbed cells inflate control-referenced correlation metrics, a phenomenon also described by Viñas Torné et al. (3). The second, *signal dilution*, occurs when error metrics dilute meaningful biological signals arising from perturbations with a small but significant number of differentially expressed genes (DEGs), causing mean predictions to appear accurate. Together, these effects allow uninformative predictors to achieve deceptively high scores even when they fail to capture underlying biology, highlighting the need to evaluate whether common benchmarking metrics can distinguish null from meaningful predictions in the first place.

Here, we address this challenge by introducing a framework for evaluating the calibration of perturbation model benchmarking metrics (Fig. 1). In this context, “calibration” refers to the degree to which a metric improves in response to a meaningful biological signal and correctly ranks informative predictions above null ones. We formalize the use of positive (informative predictor) and negative (null predictor) controls, introduce a new “interpolated duplicate” baseline that provides a stable positive control, and define a quantitative measure of calibration, the *Dynamic Range Fraction* (DRF). From an analysis of 14 Perturb-seq datasets and 13 metrics, we find that common metrics such as MSE and control-referenced Pearson correlations are often poorly calibrated, whereas weighted and rank-based alternatives such as weighted MSE (WMSE) (9), weighted 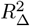 (9), and Normalized Inverse Rank (NIR) (10) exhibit consistent calibration. We then evaluated eight deep learning models across four datasets and two prediction tasks. Under well-calibrated conditions, deep learning models generally outperform uninformative baselines, indicating that the limitations reported in prior studies largely reflect shortcomings of evaluation rather than of modeling.

**Fig. 1.**
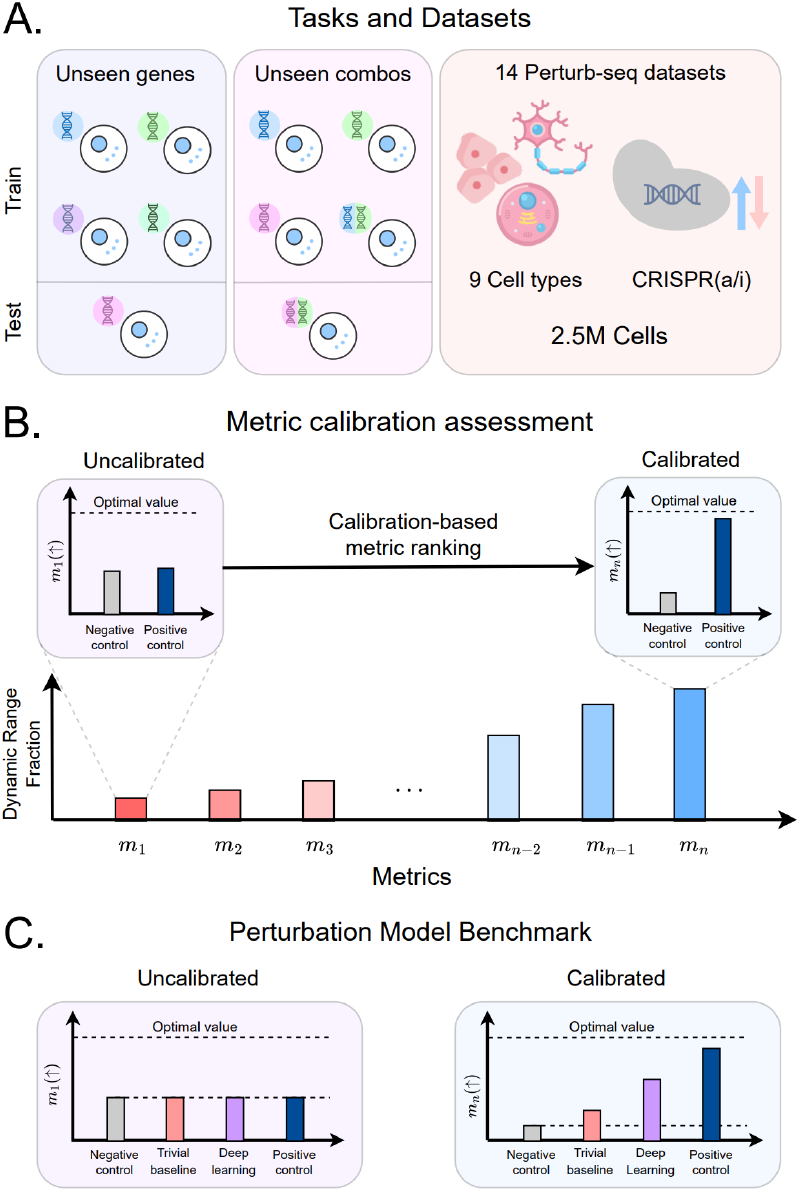
Overview of the study. (A) Benchmarking tasks: predicting unseen single-gene and gene-combination responses. We analyze 14 datasets comprising *∼*2.5 million cells across 9 cell types and multiple perturbation modalities. (B) Framework to assess metric calibration, quantified by the meta-metric *Dynamic Range Fraction* (DRF). Calibration varies widely across 13 metrics. (C) With well-calibrated metrics, positive controls outperform negative controls, and deep learning models outperform trivial baselines.

## Results

### Negative and positive controls

In biological experiments, positive and negative controls verify that an assay can detect true effects. A negative control is a condition expected to yield a null result, while a positive control is one expected to produce a consistent, strong signal. In perturbation modeling, an uninformative baseline that always predicts the mean of all perturbations (the “mean baseline”, see Sup. Note 1) serves as an effective negative control. However, benchmarks typically lack an equivalent positive control, leaving ambiguity when models perform poorly: low scores may indicate either genuine model failure or insufficient metric sensitivity to detect positive model performance.

In our previous work, we introduced the *technical duplicate* baseline as a positive control for perturbation modeling (9). This baseline splits the population of cells receiving each perturbation into two halves, using one subset to predict the other (Fig. 2A). Because both halves share the same perturbation-specific signal, the technical duplicate should, in principle, outperform null predictors. We evaluated its performance on two benchmarking datasets, Norman19 and Replogle22 K562 genome-wide Perturb-seq (GWPS), using two common metrics: mean squared error (MSE) and control-referenced delta correlation (Pearson(Δ_ctrl_)) (Fig. S1). As expected, the technical duplicate exceeded the mean baseline on both metrics in Norman19 (Fig. S1A,C). In contrast, in Replogle22 K562 GWPS it performed comparably to the mean baseline under Pearson(Δ_ctrl_) (Fig. S1D) and, unexpectedly, *underperformed* it with MSE (Fig. S1B).

**Fig. 2.**
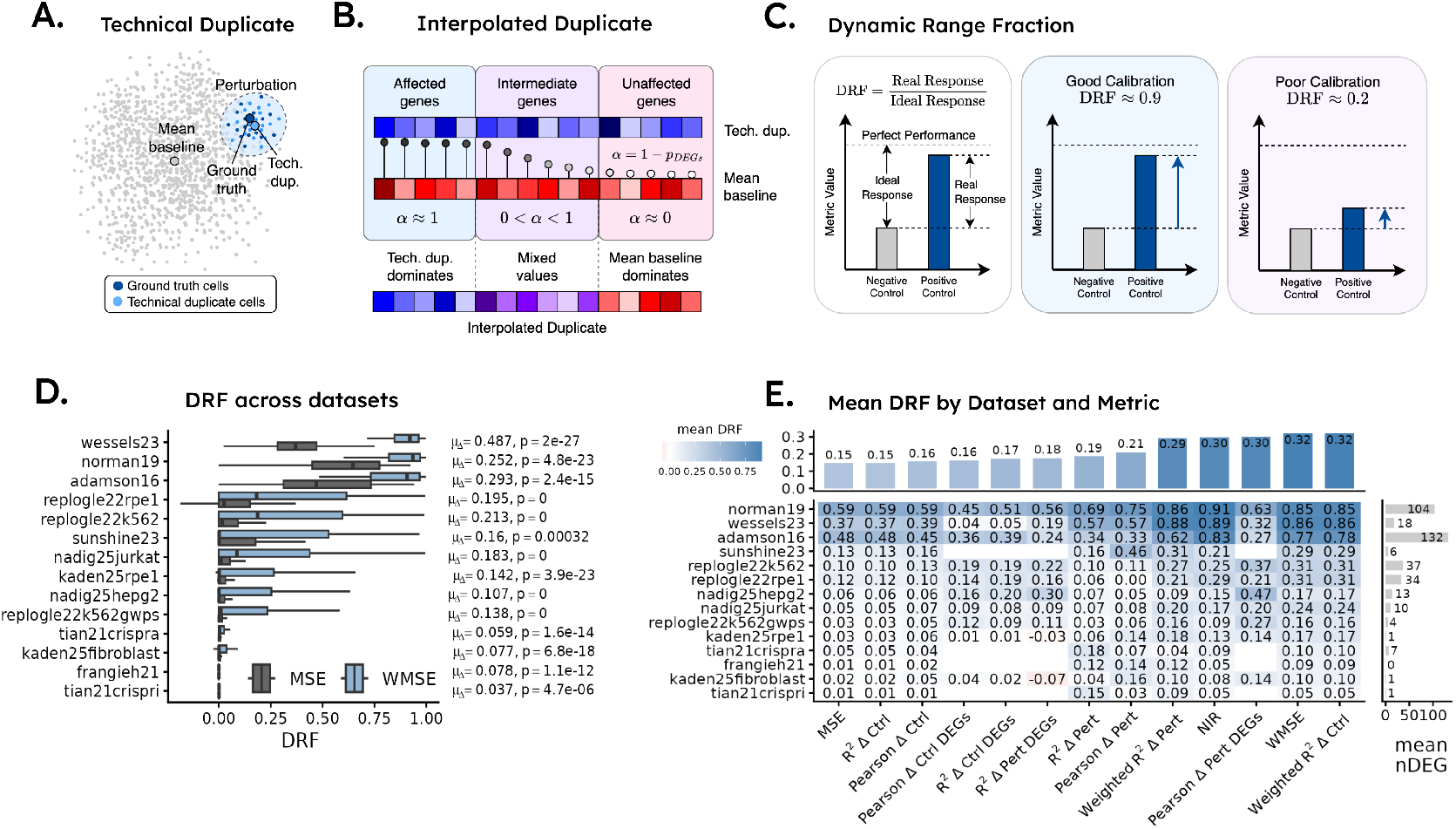
Evaluating the calibration of benchmarking metrics. (A) The *technical duplicate* baseline is generated by splitting cells from each perturbation into two groups, using one half to predict the other. (B) The *interpolated duplicate* baseline, which blends the technical duplicate and mean baseline using a weight *α* = 1 *− p*_DEGs_ from DEG analysis (adjusted p value). Strongly perturbed genes are weighted toward the technical duplicate, while weak or null genes are weighted toward the mean baseline. (C) Schematic illustrating how DRF measures calibration as the fraction of the theoretical dynamic range of a metric that is realized empirically. For each perturbation, the *ideal response* is the difference between the negative control (typically the mean baseline) and perfect performance (e.g., Pearson = 1, MSE = 0), while the *real response* is the difference between the positive control (interpolated duplicate) and the negative control. DRF is defined as the ratio of real to ideal response, representing the proportion of measurable dynamic range captured by a metric. (D) Box plots of DRF values across perturbations for MSE versus WMSE across all fourteen datasets analyzed. *µ*_Δ_ is the mean of all perturbation-wise differences between WMSE and MSE and *p* is the adjusted p-value from paired Wilcoxon rank-sum tests between WMSE and MSE. (E) Heatmap summarizing the calibration (in terms of mean DRF) of all 13 benchmarking metrics across all 14 datasets. Bar plots above and to the right of the heatmap show the mean DRF per dataset and the mean number of differentially expressed genes (DEGs) per perturbation, respectively. Cases where there are less than 10 perturbations for metric calculation (due to a low number of DEGs) in the dataset are censored.

To better understand this result, we compared MSE(technical duplicate) and MSE(mean baseline) for each perturbation in Replogle22 K562 GWPS. The technical duplicate underperformed the mean baseline for approximately 95% of perturbations (Fig. S1E). Examining representative cases of high (SUPT6H, 435 DEGs), moderate (OGFOD1, 31 DEGs), and low (COX5A, 5 DEGs) perturbation strength showed that genes strongly-affected by a perturbation (adjusted *p*_DEG_ *<* 0.05) were predicted more accurately by the technical duplicate, whereas unaffected genes (*p*_DEG_ *>* 0.95) favored the mean baseline (Fig. S2). Across all perturbations, technical duplicate performance improved with increasing perturbation strength (Fig. S1E). A downsampling titration experiment further revealed strong dependence on sampling power: increasing the number of cells used to compute the technical duplicate markedly improved its performance relative to the mean baseline (Fig. S3). Together, these findings suggest a straightforward explanation for poor technical duplicate performance: for strongly affected genes, the technical duplicate captures the true signal despite limited sampling, whereas for unaffected genes, the mean baseline provides a more stable estimate because it averages over thousands of cells to calculate unaffected gene expression (Fig. S4).

Motivated by these observations, we developed a novel *interpolated duplicate* baseline that combines the strengths of both approaches (see Sup. Note 1; Fig. 2B, Fig. S5) to create a more stable positive control. This predictor linearly interpolates between the mean baseline and the technical duplicate, weighting each gene by its DEG *p*-value (adjusted): genes with strong evidence of differential expression are shifted toward the technical duplicate, while genes with weak or null effects are shifted toward the mean baseline. This design yields a robust positive control that retains stability in weakly perturbed genes while preserving the ability to detect true perturbation effects in strongly perturbed genes (Fig. S5A,B, Fig. S6).

### Measuring the calibration of benchmark metrics

With our positive control (interpolated duplicate) and negative control (mean baseline) in hand, we next defined a *meta*-metric to quantify the calibration of benchmarking metrics. We term this quantity the “Dynamic Range Fraction” (DRF), which measures, for each perturbation, how effectively a metric distinguishes positive from negative controls. For any metric *m*(*·*), DRF is defined as

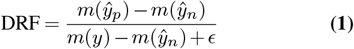

where *ŷ*_*n*_ is the mean baseline (or any other negative control), *ŷ*_*p*_ is the interpolated duplicate (or any other positive control), *y* represents the ground truth and hence yields perfect performance on the metric (for example, *MSE* = 0 or *P earson*(Δ_ctrl_) = 1), and *ϵ* is a small constant added for numerical stability. As illustrated (Fig. 2C), DRF captures the fraction of the theoretical dynamic range [*m*(*y*) *− m*(*ŷ*_*n*_)] (“ideal response”) that is realized by the empirical range [*m*(*ŷ*_*p*_) *− m*(*ŷ*_*n*_)] (“real response”). When the positive control outperforms the negative control (and hence the metric is better calibrated), DRF values increase in proportion. In a miscalibrated scenario, DRF values approach zero, reflecting limited metric sensitivity to detect improvements over the negative control. While in this study, we generally use the interpolated duplicate as the positive control and the mean baseline as the negative control, one could substitute any experimentally-appropriate controls and still utilize the DRF to measure calibration.

We applied the DRF to assess the calibration of MSE and Pearson(Δ_ctrl_) in Replogle22 K562 GWPS and the Replogle22 K562 essential-gene dataset (termed “Replogle22 K562” here). The latter targets 2,045 genes required for cell proliferation or survival and thus exhibits stronger perturbation effects (11). In both datasets, most perturbations show low DRF values, indicating poor calibration overall; Pearson(Δ_ctrl_) performs slightly better than MSE, and calibration improves with stronger perturbations (Fig. S7A).

We hypothesized that low DRF may reflect dilution of weak yet biologically meaningful signals rather than the absence of learnable structure. Indeed, for the SP2 transcription factor (TF) in Replogle22 K562 GWPS, DRF(MSE) is near zero (Fig. S7B), but gene set enrichment analysis (GSEA) of SP2 differential expression ranks recovers SP2 binding targets among the top ENCODE/ChEA gene sets (Fig. S7C). Similar trends across other transcription factors (Fig. S7D, Supplementary File 1) suggest that MSE underweights sparse but meaningful effects, a bias captured by its low DRF(MSE).

We previously proposed weighted metrics such as weighted MSE (WMSE) (9) to address the dilution of small but significant biological signals by MSE. Applying this approach largely rescues miscalibration in the Replogle22 K562 GWPS dataset, including for SP2 and the other TF perturbations we tested (Fig. S7B). Across datasets, WMSE also shows substantially better calibration than MSE, even for datasets with few DEGs (Fig. 2D, Fig. S7E).

Extending this analysis, we evaluated the calibration of 13 benchmarking metrics across 14 Perturb-seq datasets (Fig. 2E). Weighted 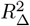, NIR, and WMSE consistently showed strong calibration across most datasets, whereas commonly used metrics such as MSE and Pearson(Δ_ctrl_) performed well only in datasets with strong perturbation signals (e.g., Norman19). When recalculating DRF using the technical duplicate as the positive control (Fig. S8), weighted 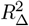, NIR, and WMSE again display consistent calibration, confirming that their apparent calibration is not an artifact of the interpolated duplicate. However, this analysis also highlights the reduced stability of the technical duplicate across metrics, further supporting the use of the interpolated duplicate as a more robust positive control.

### Benchmarking the unseen gene prediction task

Having evaluated metric calibration across our benchmarking datasets, we next assessed model performance on the unseen single gene prediction task. In this task, models are trained to predict gene expression responses for single gene perturbations that are held out during training (Fig. 3A, left). Eight models were evaluated: scGPT (12), GEARS (13), PRESAGE (14), scLambda (15), and four models that predict gene expression from foundation-model gene embeddings using a multilayer perceptron (foundation MLP or “fMLP” models), including embeddings from Geneformer (16), ESM2 (17), scGPT (12), and GenePT (18). We also included the linear baseline introduced by Ahlmann-Eltze et al. (2), which performs PCA on the training-set gene expression matrix and fits a linear model to predict unseen gene perturbation responses using these features.

**Fig. 3.**
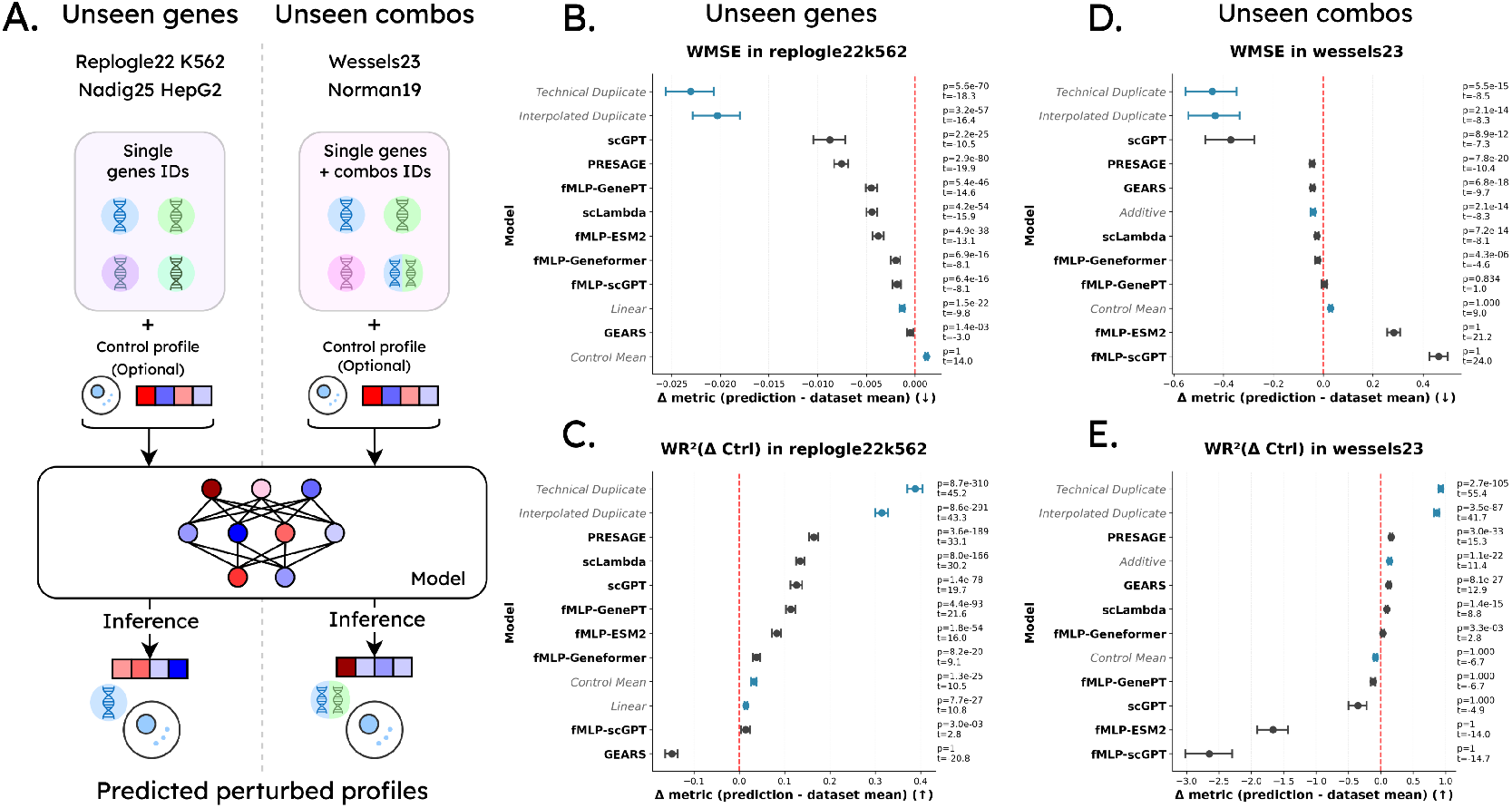
Benchmarking results under well-calibrated metrics for the Replogle22 K562 unseen-gene prediction task and the Wessels23 combination-gene prediction task. (A) Overview of benchmarking tasks. For the unseen gene prediction task, models learn to predict the expression resulting from single-gene perturbations and then generalize to predicting unseen perturbations. This task uses the Replogle22 K562 (essential genes) and Nadig25 HepG2 datasets. For the unseen gene combo task, models are shown all single genes during training along with some combo examples. They are then evaluated on predicting held-out gene combos. This task uses the Wessels23 and Norman19 datasets. (B) Forest plot showing per-perturbation performance deltas relative to the mean baseline for WMSE in Replogle22 K562, with 95% confidence intervals. Statistics from paired one-way t-test (model vs mean baseline) with Bonferroni correction. (C) Same as (B), but evaluated using the weighted 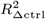 metric, where higher values indicate better performance. (D) Same as (B) but for the Wessels23 combo gene prediction dataset. (E) Same as (C) but for the Wessels23 combo gene prediction dataset.

In the Replogle22 K562 benchmarking dataset, similar to others (2, 3, 5, 6, 8), we find that models generally do not outperform the mean and linear baselines when evaluated with poorly calibrated metrics such as MSE and Pearson(Δ_ctrl_) (Fig. S9A,B; table 1). In contrast, under well-calibrated metrics such as WMSE, weighted 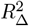, and NIR, most models outperform these baselines (Fig. 3B,C; Fig. S9C). Moreover, on all metrics, at least one deep learning model outperforms each of the linear, mean, and control baselines (table 1, table S1).

**Table 1.**
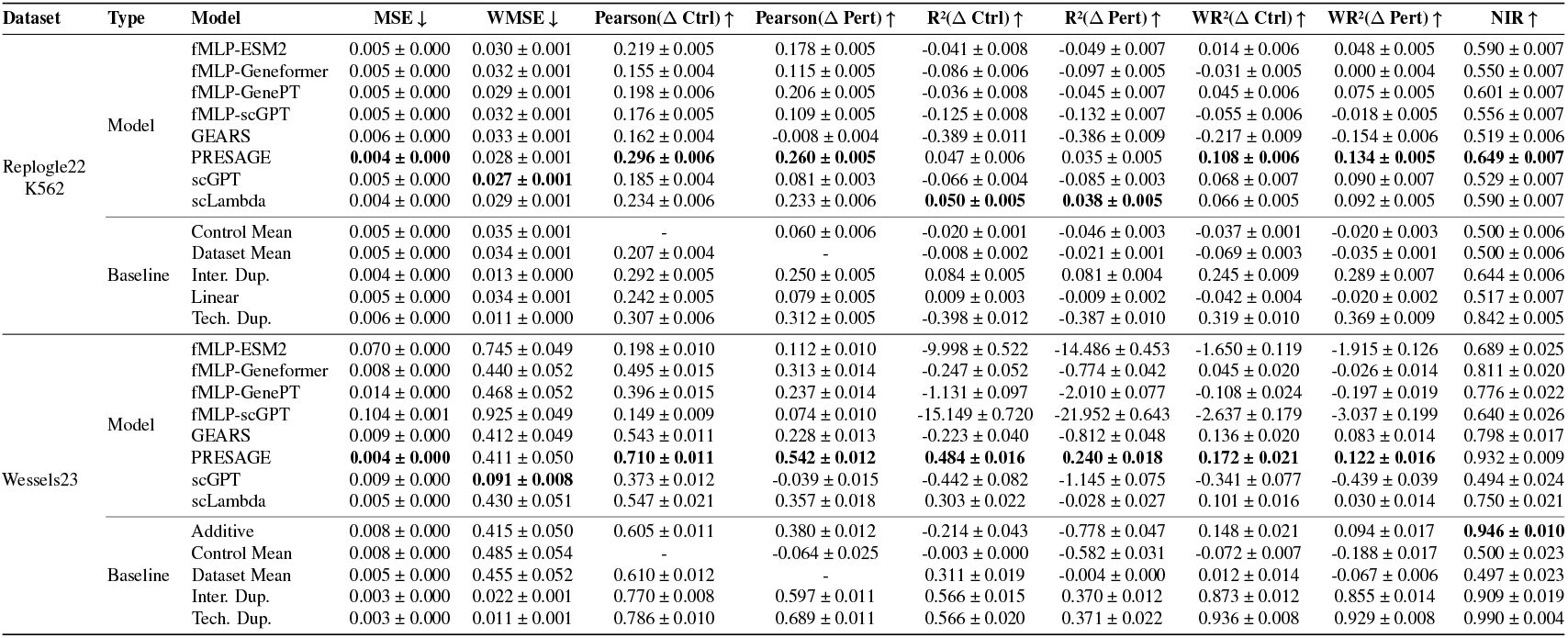
Mean performance ± standard error of the mean (SEM) for models and baselines in the unseen gene task (Replogle22 K562) and combo prediction task (Wessels23). Top-performing models / baselines are bolded (technical duplicate and interpolated duplicate excluded.) Due to space limitations, not all metrics shown (full results in table S1 and table S4).

To further validate these findings, we repeated the analysis in an independent dataset, Nadig25 HepG2 (19). The results mirrored those observed in Replogle22 K562: under well-calibrated metrics, deep learning models consistently and significantly outperformed both the mean, control, and linear baselines (Fig. S10, table S2).

Taken together, these results demonstrate that the apparent weakness of perturbation models for unseen gene prediction arises primarily from metric miscalibration rather than a lack of model performance.

### Benchmarking the unseen combos prediction task

The unseen combos prediction task evaluates whether a model can predict the effect of an unseen pair of perturbed genes after seeing both genes individually along with other gene pair examples during training (Fig. 3A, right).

We first benchmarked model performance on Norman19 (20), the most widely used dataset for this task setting (2– 6, 8). Norman19 comprises 100 single-gene and 124 double-gene perturbations. Ahlmann-Eltze et al. (2) previously compared models to a control-cell baseline and to an *additive baseline* that predicts combination effects by summing single-gene effects. The additive baseline performs strongly, which is unsurprising given that 96% of gene-pair perturbation effects are additive (2). Moreover, the combinatorial space is sparsely sampled: in the present study, the training set includes 31 of 4,950 possible pairs^2^, or 0.63%, leaving little opportunity for models to learn non-additive interactions.

Consistent with this, we found that most models exceeded the mean and control baselines across metrics, but the additive baseline remained difficult to surpass (although PRESAGE outperformed it on 5 of 13 metrics; table S3). Further analysis showed that the additive baseline captured roughly 88% of the *ideal performance gap* (see Sup. Note 1, metric calibration section) in the best-calibrated metrics, comparable to the interpolated duplicate baseline which accesses held-out combination data (Fig. S11). This evidence of additive baseline saturation, together with limited combinatorial coverage, limits Norman19’s value for benchmarking models that aim to learn non-additive genetic interactions.

To test whether models can surpass additivity when exposed to a broader range of gene combinations during training, we evaluated Wessels23 (21), which includes 28 genes and 157 unique gene pairs (train set includes 39 of 378 possible combinations, or 10.3%). In this setting, multiple models display higher average performance compared to the additive baseline under WMSE (Fig. 3D; table 1) and weighted 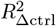 (Fig. 3E; table 1). Across all metrics, at least one model, PRESAGE, exceeded the additive baseline in 12 of 13 metrics (table 1, table S4), and consistently surpassed the mean and control baselines in well-calibrated metrics.

Similar to the unseen gene task, calibrated evaluation shows that several models consistently outperform mean and control baselines in the unseen combos task. When presented with a more robust sampling of the combinatorial search space during training (as in the Wessels23 dataset), at least one deep learning-based model (i.e., PRESAGE) beats the additive baseline in 12/13 metrics. Taken together, these findings suggest that deep learning-based perturbation models can outperform uninformative baselines on the unseen combos task, and that they may be capable of modeling non-additive gene interactions as well.

## Discussion

Our results help resolve a central tension in recent perturbation-response studies: although prior work reported that deep learning models fail to outperform uninformative baselines (2–8), we find that these reports largely reflect *metric miscalibration*, not intrinsic limits of model capacity. Common metrics such as MSE and control-referenced Pearson correlations often fail to distinguish true biological signal from noise, particularly in weakly-perturbed datasets. Using the DRF, we show that such metrics are poorly calibrated, frequently ranking null predictors above biologically meaningful positive controls. In contrast, DEG-aware and rank-based metrics (WMSE, weighted 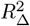, and NIR) display robust calibration across datasets, correctly rewarding models that capture perturbation-specific signals.

Under calibrated evaluation, a consistent pattern emerges across tasks: In the unseen-gene prediction setting, most deep learning-based models outperformed the mean, control, and linear baselines. This finding challenges prior reports of model failure and establishes that current architectures learn generalizable perturbation-response representations. In the combination-gene task, some models likewise exceeded null baselines and (especially for PRESAGE) even surpassed the additive baseline in the Wessels23 dataset. While models also surpassed this predictor in some metrics in the Norman19 dataset, our findings reveal the degree to which the additive baseline in Norman19 embodies saturated performance on well-calibrated metrics, and thus we recommend that future studies refrain from using this dataset as the sole benchmark of combinatorial perturbation modeling.

Additionally, PRESAGE and scLambda are notable models in our analysis, as both are newer architectures and provide novel approaches to leveraging prior knowledge for predicting perturbation effects. Based on their robust performance, it may be valuable to include these models (and potentially other newer architectures) in future benchmarking studies alongside the typically-benchmarked models like scGPT and GEARS.

Beyond our empirical findings, our study introduces further conceptual contributions. We introduce an interpolated duplicate positive control and we formalize *metric calibration* as a prerequisite for benchmarking perturbation models, analogous to assay validation in experimental biology. Without positive controls, apparent negative results may reflect low metric sensitivity rather than true model failure.

Taken together, these findings reframe the debate on the feasibility of genetic perturbation modeling. The main limitation is not model capability but the use of uncalibrated benchmarks that obscure true performance. With calibrated metrics, we show that deep learning models can exceed trivial baselines, establishing a more optimistic foundation for *in silico* perturbation modeling and its application to virtual cells for therapeutic target discovery.

### Limitations and Future Directions

Despite these advances, several limitations remain that motivate directions for future work. First, we explored only a subset of available benchmarking metrics described in prior studies. Additionally, while calibration is essential, a well-calibrated metric is not necessarily biologically meaningful, and future studies should examine which calibrated metrics best reflect interpretable and experimentally relevant perturbation responses. Second, our analysis was limited to a subset of available perturbation datasets. The comparison of performance between Norman19 (provides 0.63% of combo search space in training) and Wessels23 (provides 10.3% of search space) indicated that training set characteristics may influence the performance of both baselines and models on certain tasks. In future studies, it will be interesting to evaluate additional combinatorial datasets and more thoroughly characterize the relationship between factors like training set coverage of the combinatorial landscape and model performance. Finally, we focused on the unseen-gene and combination-gene prediction tasks, leaving the unseen-context task (predicting responses in unobserved cell types, donors, or conditions) unaddressed. Because this task directly tests the generalizability of virtual cell models, extending calibration analysis to context transfer will be a valuable extension of this work in the future.

## Dataset and Code Availability

The datasets used in this preprint were gathered from Zenodo and Figshare as described in the methods (see Sup. Note 1), and no new datasets were contributed. Code to reproduce this work is provided on Github at shiftbioscience/Perturbation-Models-Outperform-Baselines.

## ACKNOWLEDGEMENTS

We would like to acknowledge Xingyu Chen, Chloe Wang, Sebastian Schmon, and Yan Wu for their insightful feedback on this work. We would also like to thank Ricardo Henriques for providing the latex template used in our preprint.

## Supplementary Note 1: Methods

### A. Baseline definition

To evaluate metric calibration and model performance we define both uninformative and trivial baselines along with idealized positive-control baselines.

#### A.1. Control baseline

Computed at dataset level, the control baseline *µ*_*c*_ is the averaged expression profile of 8192 random unperturbed control cells.

#### A.2. Mean baseline

Computed at dataset level, it is defined as the average of averages of all train perturbations. Mathematically, given a set of *M* train perturbations each represented by a set of cells *𝒮*_1_, *𝒮* _2_, …, *𝒮*_*M*_ that can vary in size, the mean baseline is computed as:

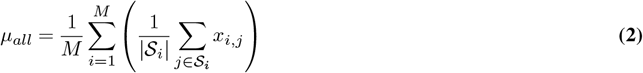

where *x*_*i,j*_ represents the expression profile of the *j*^*th*^ a cell of the *i*^*th*^ perturbation. Note that this computation is dense and weights every perturbation equally instead of every cell equally.

#### A.3. Technical duplicate

The technical duplicate baseline is defined by randomly dividing the cell set *𝒮*_*i*_ associated with perturbation *p*_*i*_ in two: a ground truth set *𝒮*_*i,GT*_, and a technical duplicate set *𝒮*_*i,T D*_. The average profiles of both sets *µ*_*i,GT*_ and *µ*_*i,T D*_ are then considered the ground truth and technical duplicate values for evaluation respectively. This definition aims to provide an idealized estimate of good performance where the prediction errors are only due to the inherent variability of the experiment. In other words, it answers the question: “*if we repeated the experiment with no batch effects, how predictive would our new experiment be?*”. Note that the technical duplicate baseline matches the sampling of the ground truth and hence has limited estimation power when the original amount of cells per perturbations is low. Additionally, to avoid any data leakage, all cells selected as technical duplicates are excluded from DEG computations to derive evaluation metric weights/masks and thus are treated as an independent duplicate data set at evaluation time.

#### A.4. Interpolated duplicate

Defined at the perturbation level, the interpolated duplicate *µ*_*i,ID*_ is a per gene interpolation between the technical duplicate prediction *µ*_*i,T D*_ and the mean baseline prediction *µ*_*all*_. Mathematically:

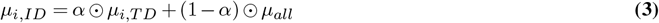

where *α* is a vector encoding DEG significance and ⊙ denotes the element-wise product operation between vectors. We define *α* = 1 *− p*_*DEGs*_ per perturbation with the help of the *p* values of an independent DEG analysis performed only on the held-out technical duplicate set. Hence, if a gene *k* is particularly affected by a perturbation, its associated interpolation value will be *α*[*k*] *≈*1. Conversely, if a gene was unaffected by a perturbation, its associated value will be *α*[*k*] *≈* 0. This setup allows the interpolated duplicate to use the predictions from the technical duplicate when there is evidence that a gene changed significantly due to a perturbation while also using the estimations of the mean baseline when the gene is unaffected.

#### A.5. Linear baseline

Defined at the perturbation level, the linear baseline is computed as in Ahlmann-Eltze et al. (2) with its original hyperparameters. In short, we perform a Principal Component Analysis (PCA) to obtain the principal component matrix *G* from the pseudobulked train dataset *Y*_train_. We then define the matrix *P* as the row selected matrix corresponding to the train perturbations and find the square matrix *W* as:

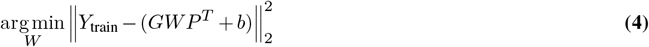

through ridge optimization with *b* the vector of row means of *Y*_train_. To perform predictions for unseen perturbations, we compute:

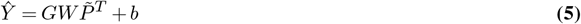

where 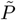is the row selected version of *G* for the desired unseen perturbations. This baseline is defined only for the unseen single perturbation task and evaluated the performance of a deliberately simple model for the task.

#### A.6. Additive baseline

Defined at the perturbation level, this baseline is computed for the unseen combination task following Ahlmann-Eltze et al. (2). For two training perturbations *µ*_*a*_ and *µ*_*b*_, their additive baseline prediction is given by:

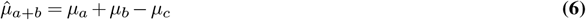

### B. Data

#### B.1. Datasets

We compiled a diverse set of 14 genetic perturbation datasets coming from 9 studies spanning 10 different cell lines under activation and repression perturbations. Detailed metadata, can be found in table S5. Datasets were obtained from the scPerturb (22) standardized data resource when available and also from the Zenodo, Figshare, or GEO records for datasets not processed by scPerturb.

#### B.2. Processing

For every dataset, we filter out any perturbation with less than 12 measured cells, any cell with less than 200 expressed genes, and any gene expressed in less than 3 cells. We use standard library size normalization (target sum of 10,000) and log1p transformation (natural logarithm) from scanpy (23). To balance the datasets, we enforce a maximum number of cells per perturbation calculated as the mean of cells per perturbation after previous filters were applied (see table S5). We further select a distinct gene set per dataset composed by the union of the top 8192 highly variable genes (scanpy’s `sc.pp.highly_variable_genes`) plus all perturbed genes.

Next, to construct our technical duplicate *𝒮*_*T D*_ and ground truth *𝒮*_*GT*_ sets, we randomly split the cells for each perturbation in two halves. While all models see the full train/val set during training (no splitting), evaluation takes place only on split data (*𝒮*_*GT*_ ). This ensures the technical/interpolated duplicate task of predicting one half of the data from the other half is mirrored in the evaluation of models as well.

Finally, we compute differentially expressed genes for every perturbation with respect to all other perturbed cells (excluding controls) using a *t −*test with overestimation of variance from scanpy’s `sc.tl.rank_genes_groups`. For each perturbation *p*_*i*_, we perform differential gene expression analysis separately for *𝒮*_*i,GT*_ and *𝒮*_*i,T D*_ against all other perturbed cells in the dataset. DEG results from the technical-duplicate half (*𝒮* _*i,T D*_) are used only for constructing the interpolated duplicate as previously mentioned. DEG results from the ground truth half (*𝒮*_*i,GT*_ ) are used only for weighted evaluation metrics such as the WMSE (9). This ensures no leakage of evaluation metric weights into the construction of the interpolated duplicate. We follow the approach of (9) computing DEGs with all other perturbed cells as the reference instead of the control population, which ensures that we evaluate over genes that make a perturbation unique and safeguards against genes changing systematically for all perturbations. The resulting processed datasets are homogeneous in terms of experimental conditions containing a single cell line and perturbation type. The only exception to the above protocol is *Replogle22k562wps* in which we randomly subsampled perturbations to 2,500 due to memory constraints.

### C. Metric calibration assessment

We numerically define metric calibration for any functional metric *m*(*ŷ*) that operates on a prediction *ŷ* and ground truth *y* with respect to positive and negative control predictions *ŷ*_*p*_ and *ŷ*_*n*_ respectively. Our proposed measure, the Dynamic Range Fraction (DRF), evaluates how much of the ideal performance gap between the perfect prediction and a negative control *m*(*y*) *−m*(*ŷ*_*n*_) is covered by the real performance gap between the positive control and negative control *m*(*ŷ*_*p*_) *−m*(*ŷ*_*n*_). Mathematically:

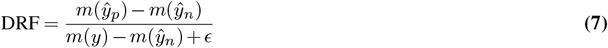

with *ϵ* being a small value for numerical stability. Note that the DRF is defined for every perturbation individually which allows the study of different factors influencing calibration. For this work, we choose the following ground truths and controls when evaluating metric calibration:

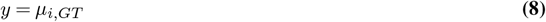

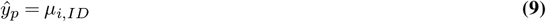

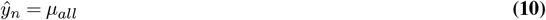

### D. Metrics

We assessed calibration and model performance on 13 metrics. At the individual perturbation level, we measured direct error using mean squared error (MSE) and its DEG weighted variant (WMSE) as proposed by Mejia et al. (9). We also assessed relative error using a suite of 10 delta-based metrics, which compare the predicted versus true expression shift from a baseline. These were generated by combining choices for three components: (i) the similarity function (Pearson correlation or the coefficient of determination *R*^2^), (ii) the reference baseline (the mean of the control population *µ*_*c*_, or the perturbed mean baseline *µ*_*all*_), and (iii) the gene set (all genes, only DEGs, or a continuous weighting scheme for *R*^2^ as proposed in Mejia et al. (9)). To evaluate the global structure of the prediction space, we also used the Normalized Inverse Rank (NIR) introduced by Wu et al. (10) (described as the “transposed-rank” metric in that work). This metric ranks the Euclidean distance of a prediction to its correct ground truth against its distances to all other ground truth perturbations.

### E. GSEA analysis

We performed a gene set enrichment analysis using the fgsea package (v1.32.4), with gene-set `minSize = 10` and `maxSize = 5000`, on overlapping perturbations from the K562 genome-wide perturbation dataset and the ENCODE and ChEA consensus TFs from ChIP-X library (24) (*n* = 90 perturbations). We used the *t −* statistics from our differential gene expression analyses performed on one-half of each perturbation vs the rest of the perturbation space as scores. For each TF, we ranked all the gene sets tested using absolute normalized enrichment scores (NES) to determine the rank of the self-enrichment signals. We selected compelling examples with low number of significant DEGs and low rank as evidence of strong biological signal. All analyses were performed in R v4.4.1.

### F. Benchmarked deep learning models

Open-source models benchmarked includes scGPT (v0.2.4) (12), GEARS (v0.1.2) (13), and scLambda (not versioned) (15) using default parameters. Additionally, we benchmarked PRESAGE^3^ (not versioned) (14) using default parameters. All models were implemented as close as possible to their original version by wrapping the source codebase in a runner script which was executed in model-specific docker containers. The only notable modifications to official releases are the replacement of `nn.Embedding` normalization by built-in normalization in the GeneEncoder class of scGPT (as recommended by the authors), and the adaptation of PRESAGE^4^ to predict combos by adding the latent embeddings of combo genes before the pooling operation.

In addition to models designed for perturbation prediction, we also evaluated the utility of gene embeddings extracted from foundation models with an MLP probing approach. To this end, we collected gene representations from Geneformer (16), GenePT (18), ESM2 (17) and scGPT (12), and finetuned a 6-layer fully connected network to predict gene expression profiles from these embeddings. The hidden dimension was set to 512 and the latent dimension in the bottleneck layer 3 was set to 128 with a leakyReLU activation function. For the unseen single perturbation task, we trained with single gene embeddings as input while for the combination task we added gene the embeddings’ encoded representation in the bottleneck layer and then decode back to expression space. In all datasets, we train for 100 epochs with a batch size of 256 and a learning rate of 0.001 for the Adam optimizer (25) minimizing an MSE loss.

### G. Benchmark setup

For the *unseen single perturbation* task, we perform 5-fold cross-validation, using a 70/10/20% train/validation/test split in each fold. For the *unseen combo perturbation* task, we perform 2-fold cross-validation over the set of combination perturbations. Specifically, all single perturbations are retained in the training set, while double perturbations are randomly divided into two equal halves. In each fold, one half is used for training and validation and the other for testing. Within the training half, combinations are further split evenly so that 25% of all combinations are used for training and 25% for validation.

As previously noted, cells that belong to the technical duplicate set *S*_*T D*_ are used for training if their perturbation is assigned to the training set. However, when computing evaluation on the test set, only the ground truth cells in *S*_*GT*_ are used as true values and the technical duplicate cells in *S*_*T D*_ are used as held out independent predictions for baseline computation.

### H. Statistical tests

When comparing metric results across conditions, comparisons were made using statistical tests paired by perturbation. For parametric comparisons, a one-way dependent Student t-test was performed. T-statistics and Bonferroni corrected p-values were reported. For non-parametric comparisons, a paired one-sided Wilcoxon signed rank test was performed. The average difference of values (perturbed value - negative control value) was reported as *µ*_Δ_ along with corrected p-values. For forest plots, we estimated 95% confidence intervals via bootstrapping (sampling 10,000 times).

## Supplementary Figures

**Fig. S1.**
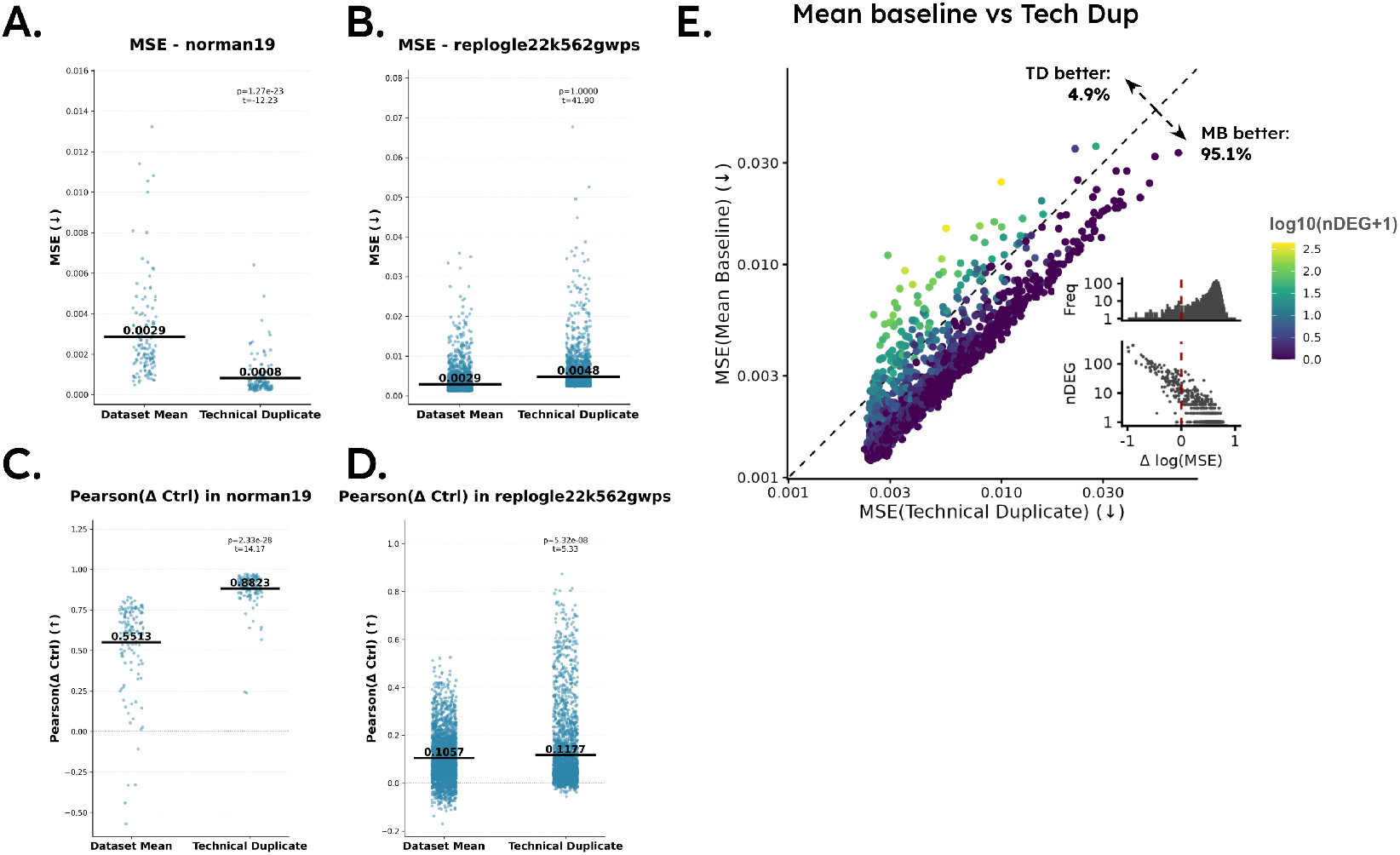
Performance of dataset mean and technical duplicate baselines across datasets and metrics. (A) Strip plots comparing the mean baseline and technical duplicate baseline by mean squared error (MSE) against ground-truth expression in the Norman19 dataset. (B) Same as (A), but for the Replogle22 genome-wide Perturb-seq (GWPS) dataset. (C–D) Same as (A–B), evaluated using the control-referenced Pearson correlation (Pearson(Δ_ctrl_)) instead of MSE. (E) For each perturbation in the Replogle22 GWPS dataset, comparison of errors from the technical duplicate versus the mean baseline, with points colored by the number of significant differentially expressed genes (DEGs). Inset plots show the distribution of the differences between technical duplicate and mean baseline MSE levels along with the relationship between these deltas and the number of DEGs in the perturbation.

**Fig. S2.**
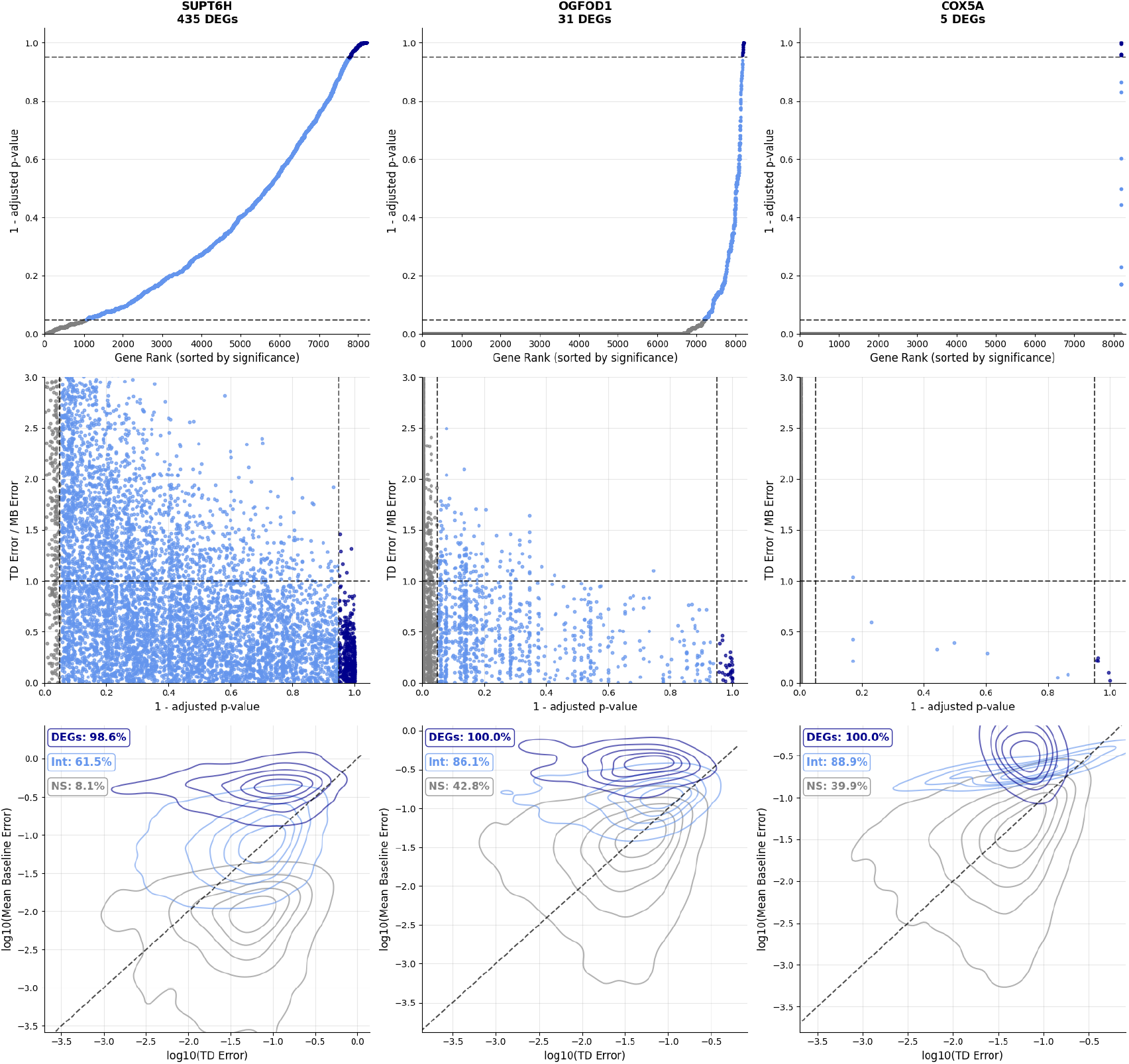
Relationship between differential gene expression and technical duplicate performance in the Replogle22 K562 GWPS dataset. Top row: rank plots showing the distribution of genome-wide adjusted *p*-values from differential expression analysis for three representative perturbations (SUPT6H, OGFOD1, and COX5A). Middle row: scatter plots relating gene-wise adjusted *p*-values to the ratio of technical-duplicate (TD) error (vs. ground truth) to mean-baseline (MB) error. Bottom row: contour plots comparing technical-duplicate and mean-baseline errors, with genes grouped as affected (adjusted *p <* 0.05), unaffected (*p >* 0.95), or intermediate (0.05 *< p <* 0.95). Percentages denote the fraction of genes where the technical duplicate outperforms the mean baseline as a predictor.

**Fig. S3.**
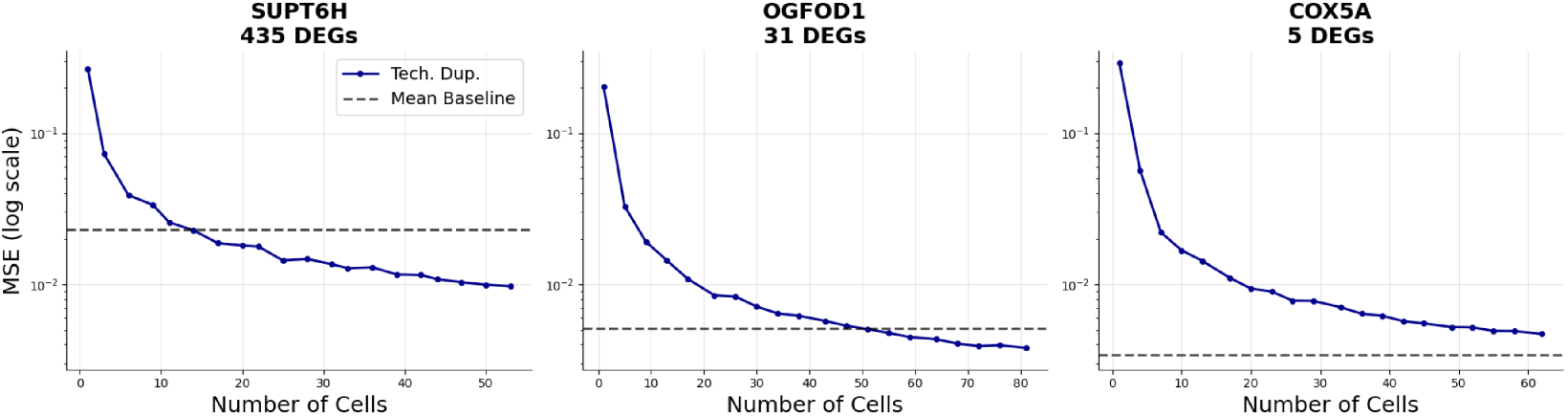
Effect of sample size on technical-duplicate performance in the Replogle22 K562 GWPS dataset. Line plots show the relationship between the number of cells sampled per perturbation and mean squared error (MSE) against ground truth for three representative perturbations. The mean-baseline performance is shown in each case as a reference.

**Fig. S4.**
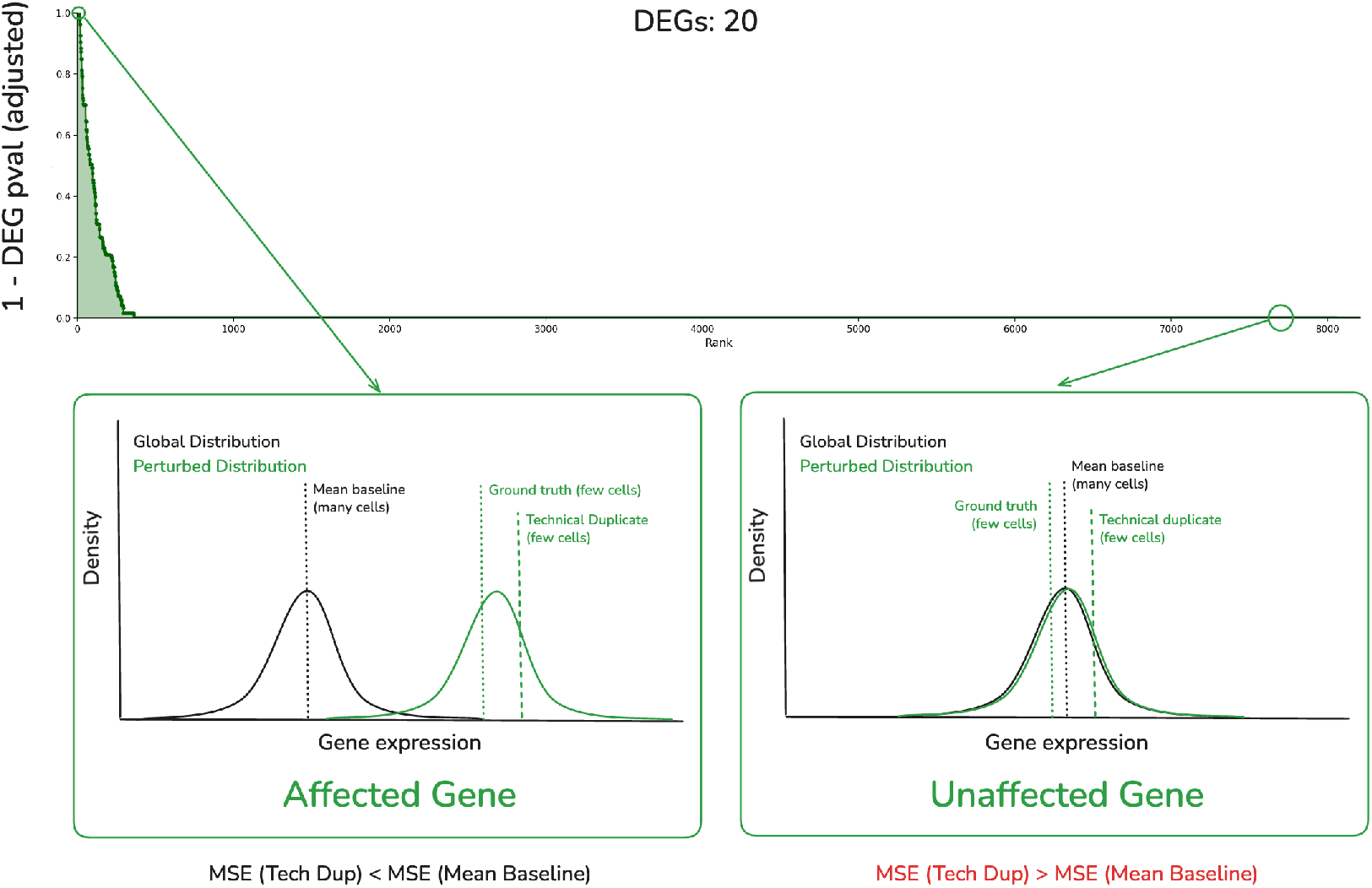
Conceptual illustration of factors influencing technical-duplicate performance relative to the mean baseline. For genes strongly affected by perturbation, the true expression distribution shifts substantially, allowing the technical duplicate to predict the ground truth more accurately despite limited cell numbers. For weakly or unaffected genes, the mean baseline provides a more stable estimate of unperturbed expression, as the sampling noise in the technical duplicate exceeds the true perturbation effect, leading to poorer predictive performance.

**Fig. S5.**
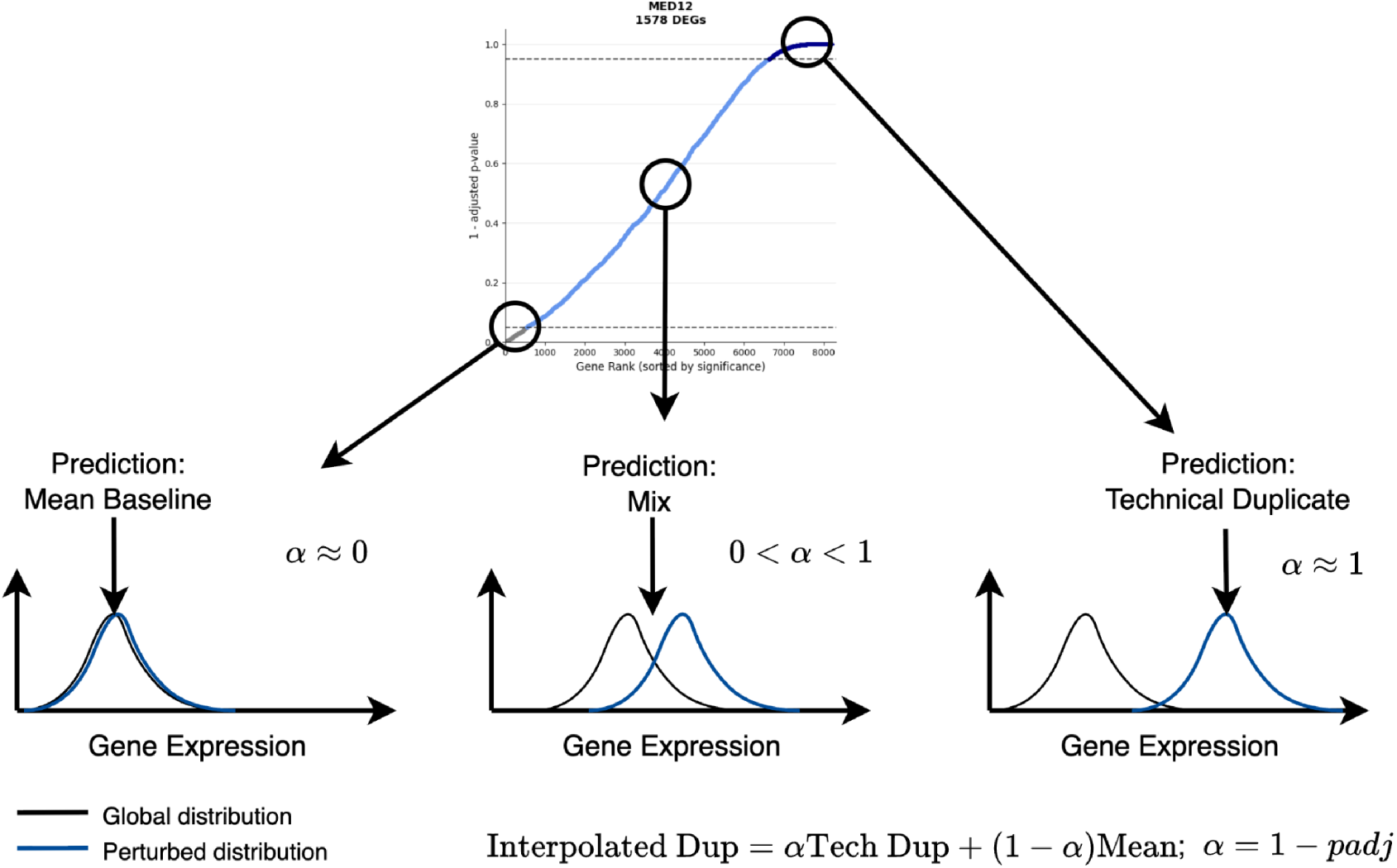
Schematic illustrating the construction of the interpolated duplicate baseline. To overcome the limitations of the technical duplicate for weak perturbations, the interpolated duplicate combines the technical duplicate and mean baseline via linear interpolation based on DEG strength (see Sup. Note 1 for full definition). When a perturbation is weak (*α ≈* 0), predictions approximate the mean baseline; when strong (*α ≈* 1), predictions converge toward the technical duplicate. Most perturbations fall between these extremes, yielding a balanced mixture of the two baselines.

**Fig. S6.**
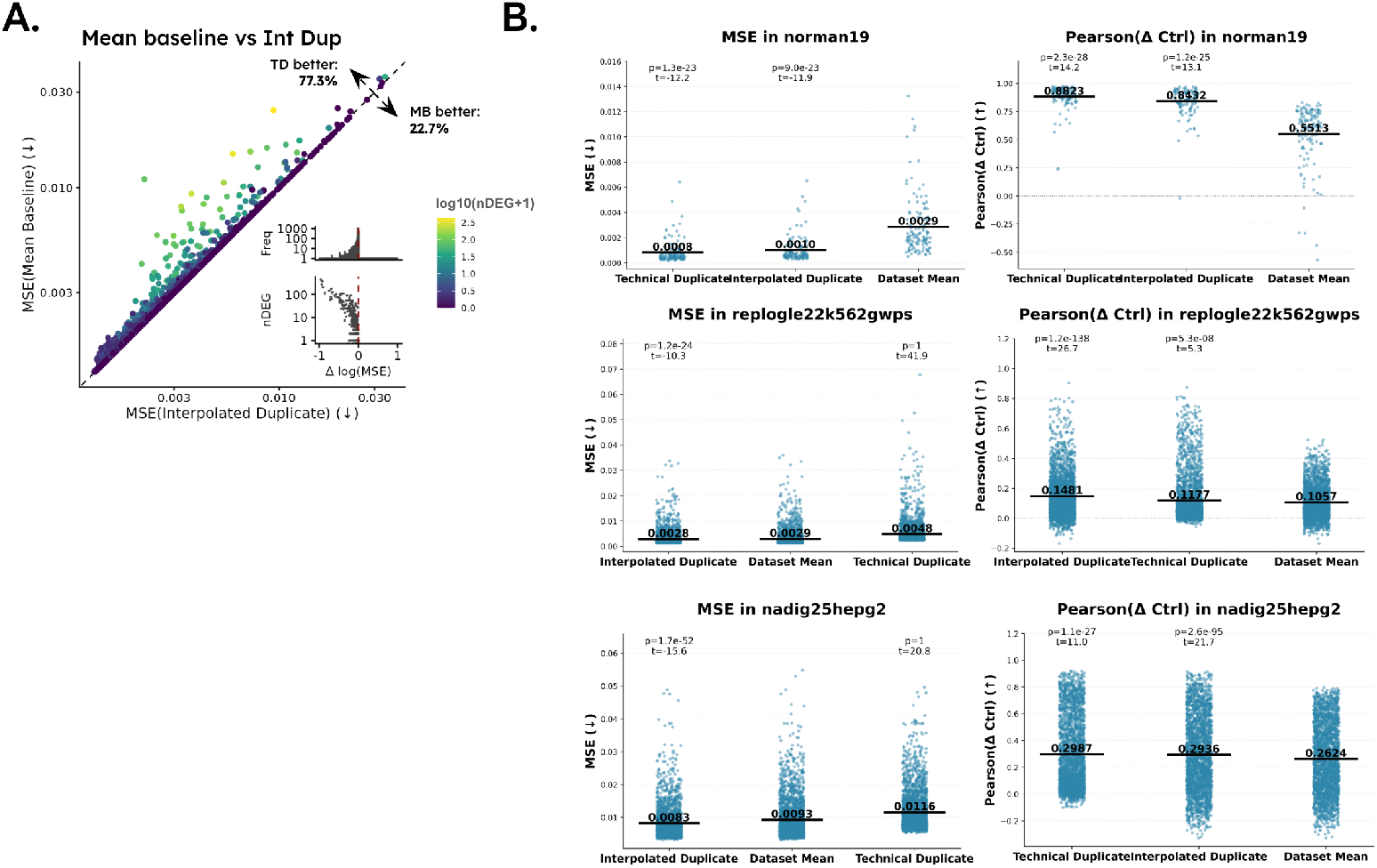
(A) For each perturbation in the Replogle22 GWPS dataset, comparison of errors from the interpolated duplicate versus the mean baseline, with points colored by the number of significant differentially expressed genes (DEGs). Inset plots show the distribution of the differences between technical duplicate and mean baseline MSE levels and the relationship between these deltas and the number of DEGs in the perturbation. (B) Strip charts showing the performance of the mean baseline, technical duplicate, and interpolated duplicate in MSE and Pearson(Δ Ctrl) metrics in three datasets (Norman19, Nadig25 HepG2, and Replogle22 K562 GWPS). Population means are reported along with *t* statistics and p-values from paired one-sided *t−*tests for the technical duplicate and interpolated duplicate vs the mean baseline. In each plot, baselines are ranked by best average performance.

**Fig. S7.**
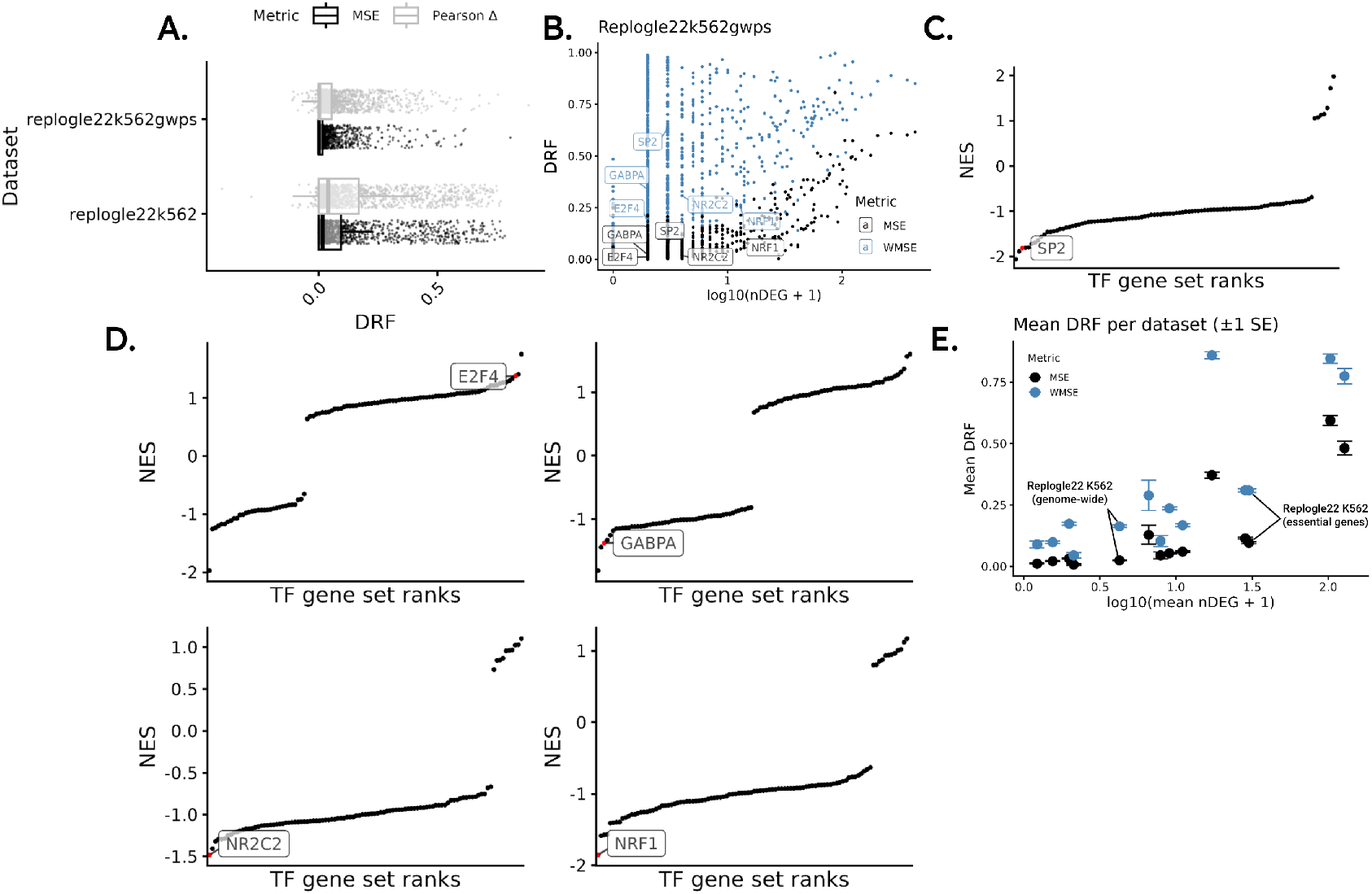
Calibration analysis and biological interpretation of metric weighting. (A) Box plots comparing Dynamic Range Fraction (DRF) values for MSE and control-referenced Pearson correlation (Pearson(Δ_ctrl_)) across perturbations in the Replogle22 K562 genome-wide Perturb-seq (GWPS) and essential-gene datasets. (B) Scatter plot comparing perturbation-wise DRF for MSE and WMSE with the number of DEGs per perturbation in the Replogle22 K562 GWPS dataset. (C, D) Ranked Normalised Enrichment Scores (NES) from a gene set enrichment analysis (GSEA) of DGE *t−*statistics (from *t*-test vs all other perturbations) for 5 perturbations, performed against the ChEA transcription factor ChIP-seq database. We labeled the perturbation position in its corresponding ChEA gene-set. (E) Scatter plot of mean DRF(MSE) and DRF(WMSE) *±*1 standard error across perturbations in each dataset showing the relationship between perturbation strength and calibration for MSE and WMSE.

**Fig. S8.**
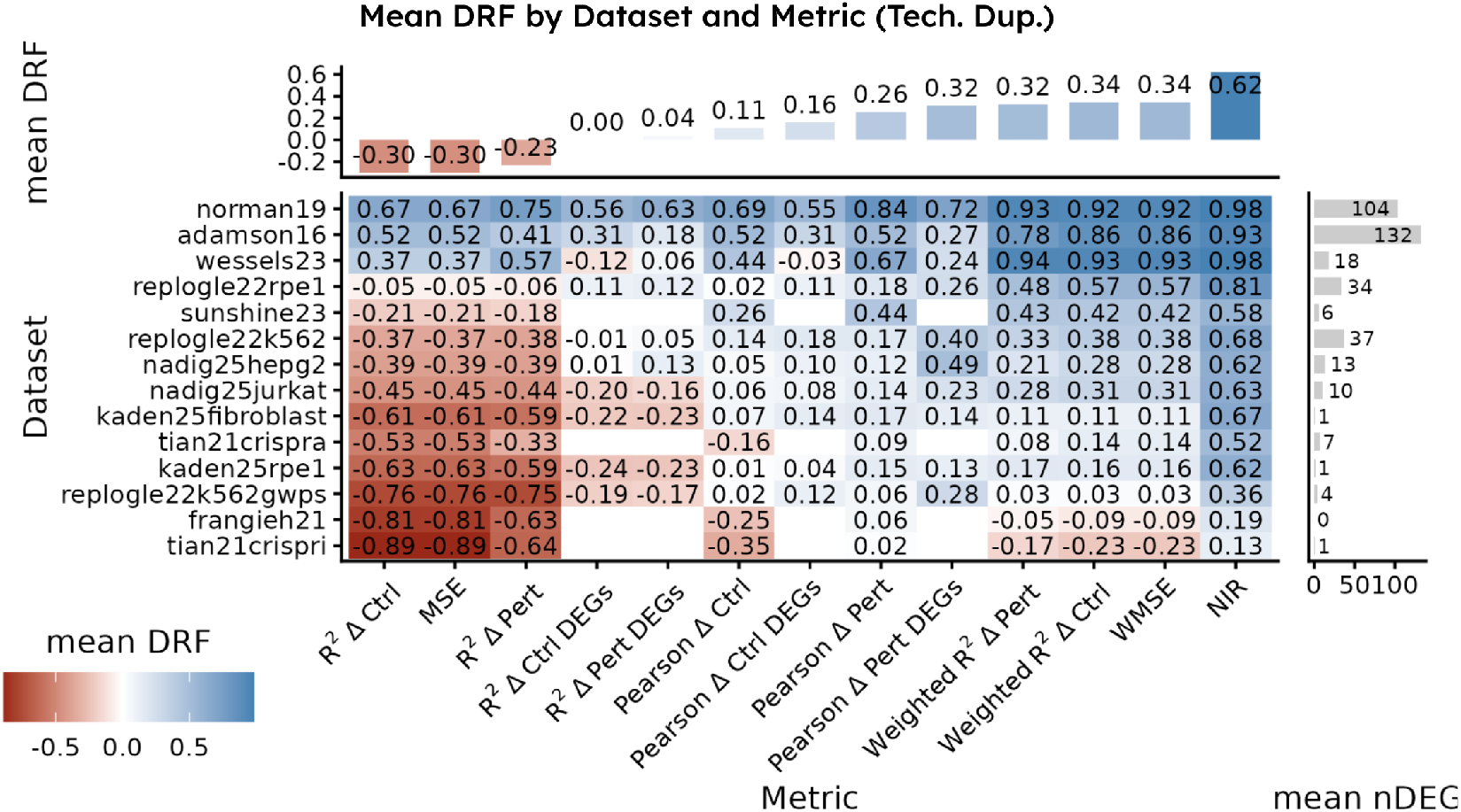
Heatmap summarizing the calibration of all 13 benchmarking metrics across all 14 datasets. DRF was calculated with the *technical duplicate* as the positive control, rather than the interpolated duplicate. Bar plots above and to the right of the heatmap show the mean DRF per dataset and the mean number of differentially expressed genes (DEGs) per perturbation, respectively. In all cases where cells have missing values, there were not enough data points after filtering for DEGs to pass the minimum threshold (*n ≥* 10) for metric calculation.

**Fig. S9.**
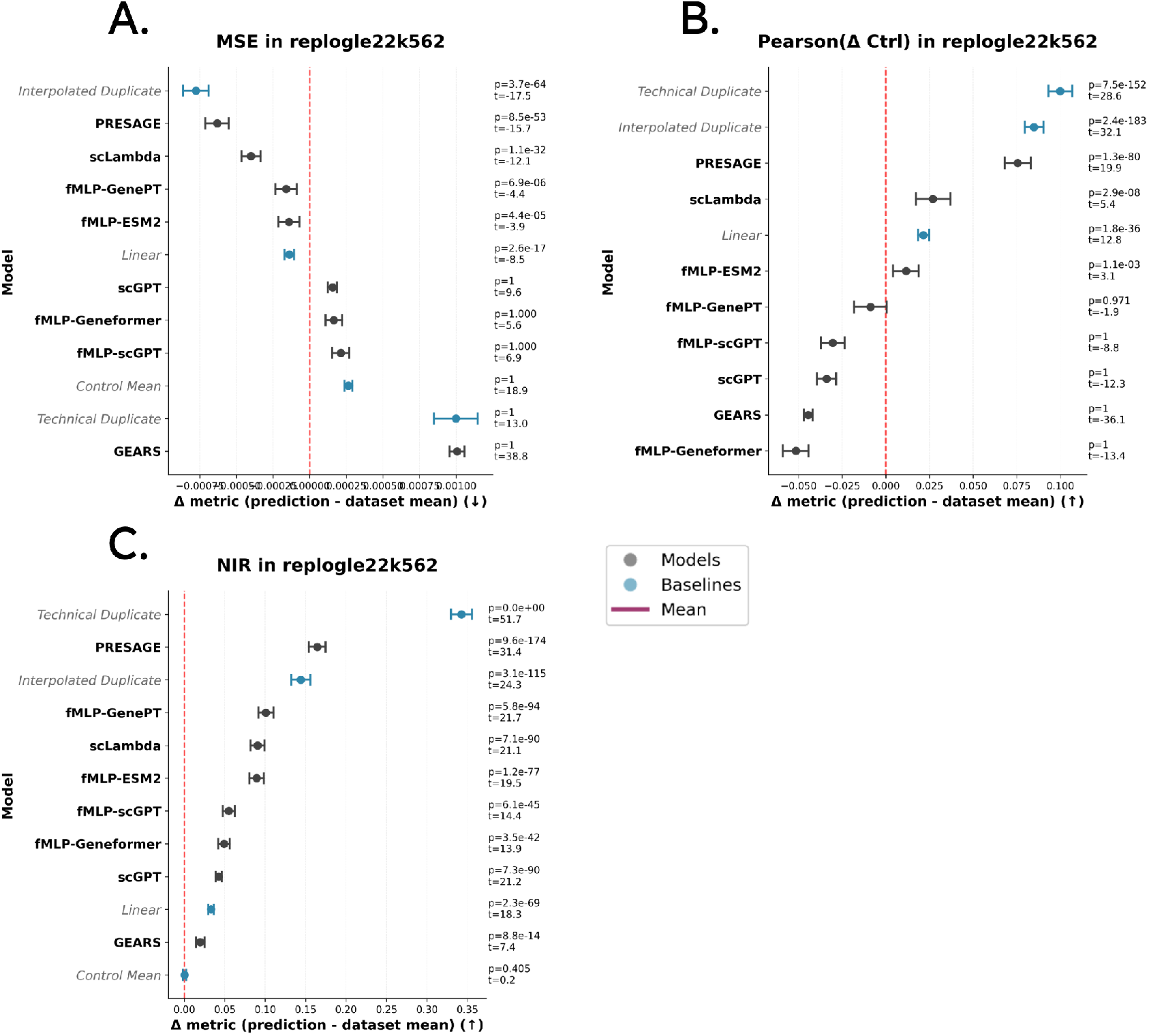
Benchmarking results under standard and well-calibrated metrics for the Replogle22 K562 unseen-gene prediction task. (A) Forest plot of per-perturbation performance differences relative to the mean baseline on the MSE metric, with 95% confidence intervals. One-sided paired *t*-tests were performed between each model or baseline and the mean baseline, Bonferroni-adjusted *p*-values and *t*-statistics are shown. (B) Same as (A), but evaluated using the control-referenced Pearson correlation (Pearson(Δ_ctrl_)), where higher values indicate better performance. (C) Same as (A) but for the normalized inverse rank (NIR) metric.

**Fig. S10.**
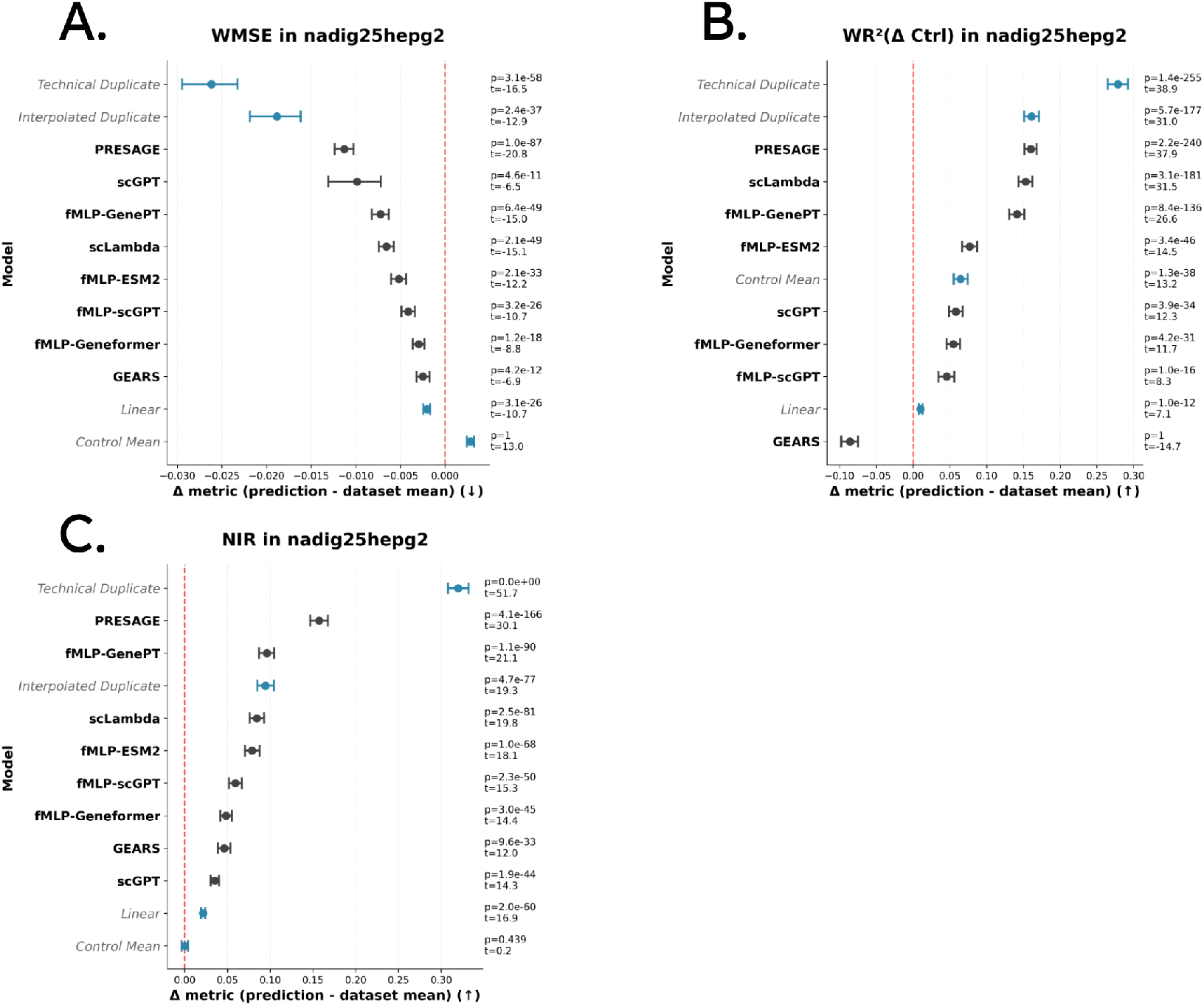
Benchmarking results under standard and well-calibrated metrics for the Nadig25 HepG2 unseen-gene prediction task. (A) Forest plot of per-perturbation performance differences relative to the mean baseline on the WMSE metric, with 95% confidence intervals. One-sided paired *t*-tests were performed between each model or baseline and the mean baseline, Bonferroni-adjusted *p*-values and *t*-statistics are shown. (B) Same as (A), but evaluated using the control-referenced weighted 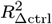 metric, where higher values indicate better performance. (C) Same as (A) but for the normalized inverse rank (NIR) metric.

**Fig. S11.**
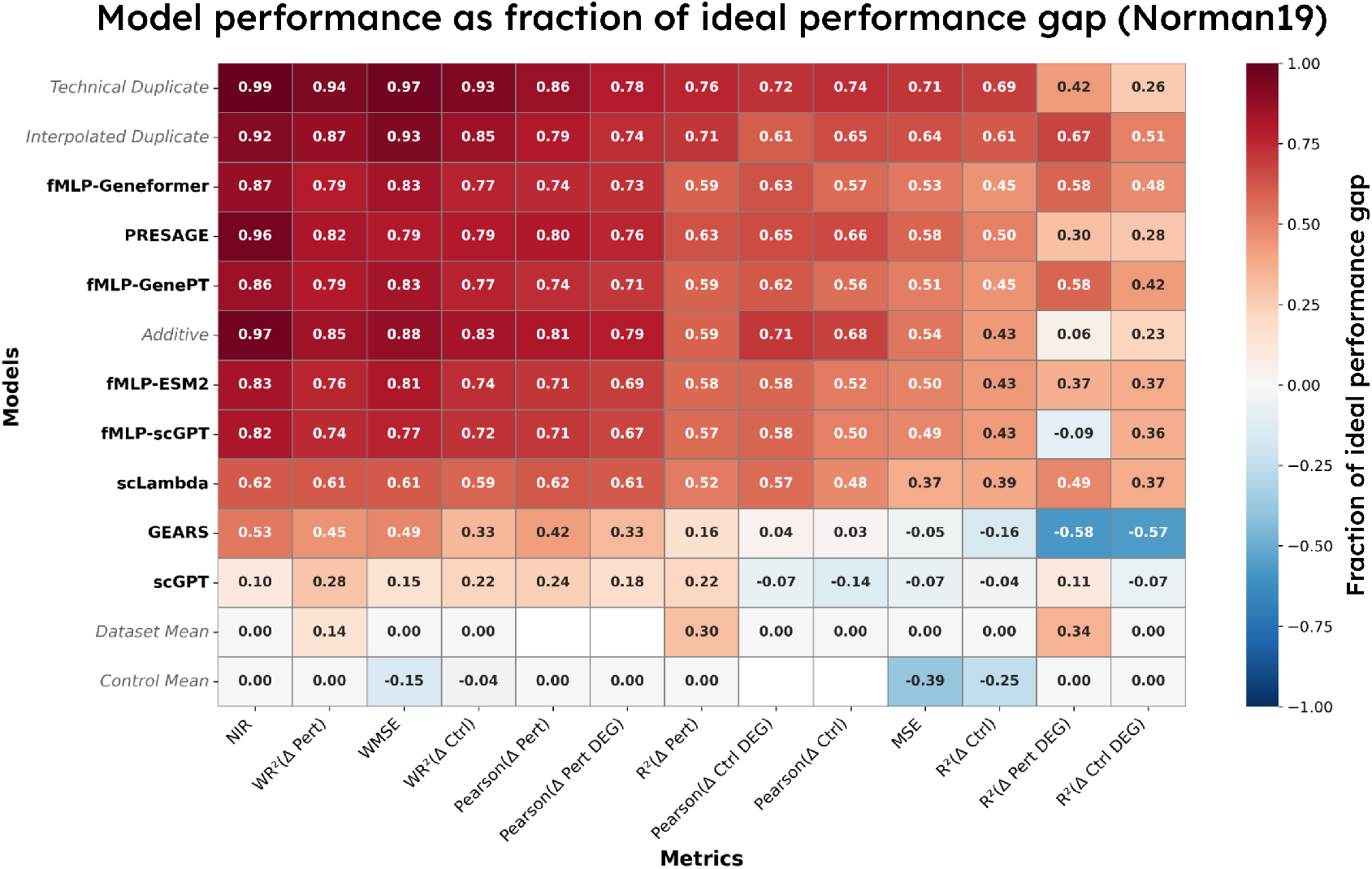
Model performance as a fraction of the *ideal performance gap* (see definition in Sup. Note 1). Briefly, heatmap displays the average performance of each model scaled between the performance of the mean baseline (or control baseline, in the case of Δ pert metrics) and the perfect score for that metric (e.g., 1 for Pearson and 0 for MSE).

## Supplementary Tables

**Table S1.**
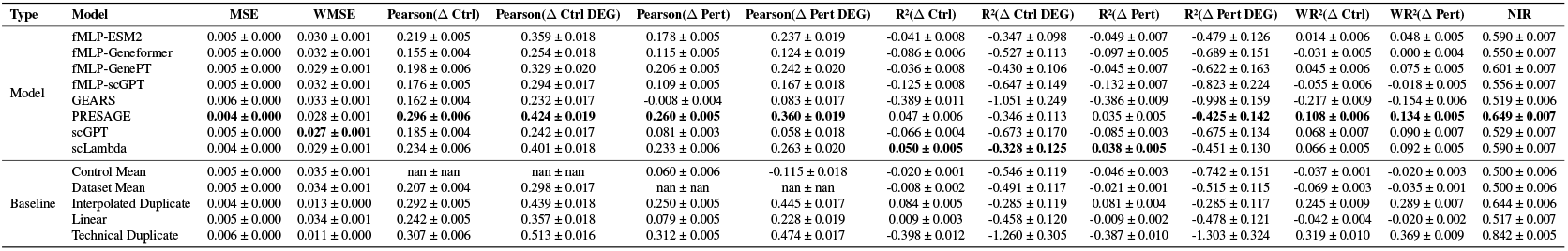
Mean performance ± SEM for Replogle22 K562 unseen gene prediction task (all metrics).

**Table S2.**
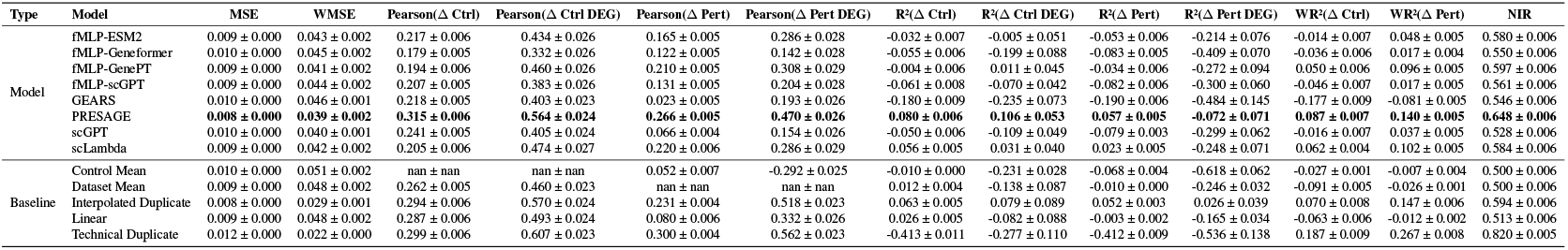
Mean performance ± SEM for Nadig25 HepG2 unseen gene prediction task (all metrics).

**Table S3.**
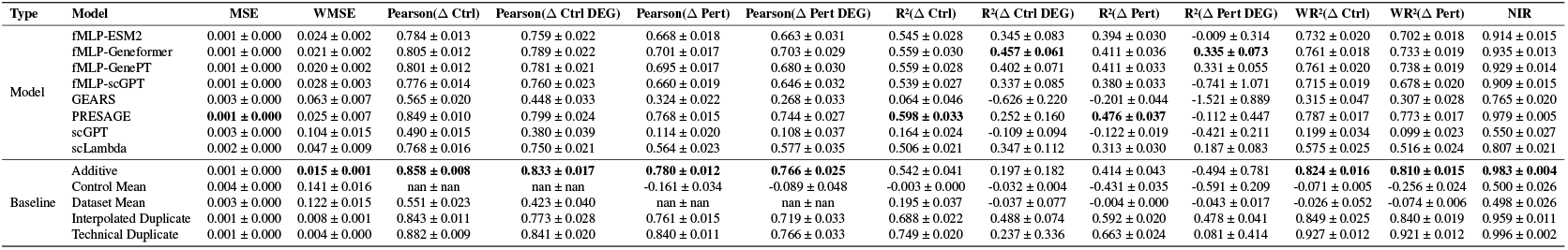
Mean performance ± SEM for Norman19 combo gene prediction task (all metrics).

**Table S4.**
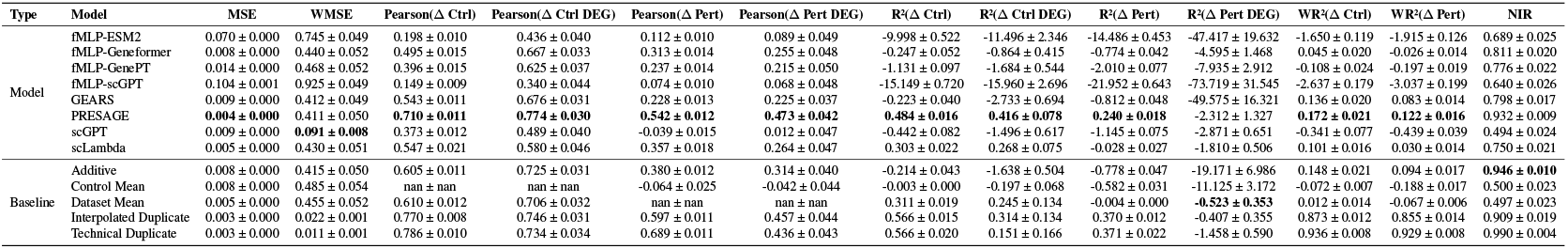
Mean performance ± SEM for Wessels23 combo gene prediction task (all metrics).

**Table S5.** Overview of single-cell perturbation datasets used in this study. Reported numbers of genes, cells, and perturbations from fully preprocessed dataset.

| Dataset | Cells | Genes | Perturbations | Cell type | Modality | CRISPR type |
| --- | --- | --- | --- | --- | --- | --- |
| adamson16 | 55,021 | 8,247 | 91 | K562 | Perturb-seq | CRISPRi |
| frangieh21 | 47,571 | 8,319 | 241 | Melanoma | Perturb-CITE-seq | CRISPR-Cas9 |
| kaden25fibroblast | 373,328 | 9,419 | 1,837 | Fibroblast | Perturb-seq | CRISPRa |
| kaden25rpe1 | 697,787 | 9,474 | 1,837 | RPE1 | Perturb-seq | CRISPRa |
| nadig25hepg2 | 109,183 | 8,746 | 2,323 | HepG2 | Perturb-seq | CRISPRi |
| nadig25jurkat | 192,708 | 8,473 | 2,347 | Jurkat | Perturb-seq | CRISPRi |
| norman19 | 88,628 | 8,221 | 238 | K562 | Perturb-seq | CRISPRa |
| replogle22k562 | 237,024 | 8,370 | 2,045 | K562 | Perturb-seq | CRISPRi |
| replogle22k562gwps | 420,765 | 8,206 | 2,492 | K562 | Perturb-seq | CRISPRi |
| replogle22rpe1 | 173,539 | 8,438 | 2,330 | RPE1 | Perturb-seq | CRISPRi |
| sunshine23 | 31,441 | 8,321 | 213 | Calu3 | Perturb-seq | CRISPRi |
| tian21crispra | 17,910 | 8,229 | 101 | iPSC-derived neurons | CROP-seq | CRISPRa |
| tian21crispri | 28,967 | 8,322 | 185 | iPSC-derived neurons | CROP-seq | CRISPRi |
| wessels23 | 22,794 | 8,210 | 187 | Monocytes | CaRPool-seq | CRISPRi |

## Footnotes

1 “Virtual cells” may also refer to multi-scale, multi-modal, and/or physicsbased in silico models of living cells and tissues (1).

2 These 31 pairs represent half of the total 62 combinations in our train/validation split. In contrast, Ahlmann-Eltze et al. (2) used all 62 combinations for training since their benchmark did not include a validation set.

3 Importantly, PRESAGE was used solely for academic benchmarking under the Genentech Non-Commercial Software License (Version 1.0, 2022). The model was not used in any discovery, pre-clinical, or therapeutic development activities. All results are reported for non-commercial academic research purposes only.

4 Our scripts which adapt PRESAGE for benchmarking are included in the GitHub repo accompanying this work, along with the original PRESAGE LICENSE and NOTICE files. We also include a MODIFICATIONS.md file specifying the changes we added. Updated or added scripts retain the original copyright notice in comment format.

